# Early language experience modulates the cortical tracking of speech

**DOI:** 10.1101/2023.09.14.557701

**Authors:** Jose Pérez-Navarro, Anastasia Klimovich-Gray, Mikel Lizarazu, Giorgio Piazza, Nicola Molinaro, Marie Lallier

## Abstract

Cortical tracking of speech is relevant for the development of speech perception skills. However, no study to date has explored whether and how cortical tracking of speech is shaped by accumulated language experience, the central question of this study. In 35 bilingual children (6 y.o.) with considerably bigger experience in one language, we collected electroencephalography data while they listened to continuous speech in their two languages. Cortical tracking of speech was assessed at acoustic-temporal and lexico-semantic levels. Children showed more robust acoustic-temporal tracking in the least experienced language, and more sensitive cortical tracking of semantic information in the most experienced language. Additionally, and only for the most experienced language, acoustic-temporal tracking was specifically linked to phonological abilities, and lexico-semantic tracking to vocabulary knowledge. Our results indicate that accumulated linguistic experience is a relevant maturational factor for the cortical tracking of speech at different levels during early language acquisition.

## Introduction

Continuous exposure to spoken language in a wide variety of contexts makes its acquisition appear spontaneous and effortless. Despite this apparent ease, developmental evidence shows that complex brain systems and processes supporting language comprehension emerge between perinatal stages and around three years of age, and become language-selective between birth and the start of primary school (for reviews see Gervain, 2015; Kuhl, 2004; Skeide & Friederici, 2016). However, little is known about how these neurocognitive processes evolve and mature as a function of language experience, an established contributor to language development (e.g., Carbajal & Peperkamp, 2020; Gámez et al., 2019; Gathercole & Thomas, 2005; Paradis et al., 2016; Pearson et al., 1997; Thordardottir, 2011).

Accurate multidimensional models of language development require understanding how different levels of language exposure and proficiency modulate fundamental brain mechanisms underlying language acquisition, including speech comprehension. We focus on the cortical tracking of speech (CTS) (Giraud & Poeppel, 2012), a neurocognitive process critical for understanding speech as it unfolds over time. CTS typically refers to the dynamic alignment of brain oscillatory activity to the temporal modulations of the speech envelope (Obleser & Kayser, 2019). In this study we refer to such speech envelope tracking as *acoustic-temporal* CTS, which takes place within delta (aligning with prosodic phrasing) and theta (syllable timing) frequency bands and has shown to support speech comprehension in adults (e.g., Ding & Simon, 2012; Gross et al., 2013; Luo & Poeppel, 2007; Molinaro & Lizarazu, 2018; Peelle et al., 2013). Beyond the speech envelope, a growing number of studies are investigating CTS at higher order lexical and semantic levels (Brodbeck et al., 2022; Broderick et al., 2018, 2021). Here, we term this process as *lexico-semantic* CTS. We hypothesize that early language experience shapes the maturation of CTS at both acoustic-temporal and lexico-semantic levels. In fact, behavioral evidence shows that language experience supports increasingly efficient processing and comprehension skills for continuous speech. For example, in monolingual environments, the amount of child-directed input is positively linked to word processing speed, the ability to encode lexical information from continuous speech input (Hurtado et al., 2008; Weisleder & Fernald, 2013). In addition, in bilingual contexts, the proportional exposure to both languages correlates with language-specific word processing speed (Hurtado et al., 2014).

We propose that language experience contributes to CTS at two different levels. First, accumulated input in a language should be crucial for building up knowledge about the temporal statistics embedded in the speech envelope, which contributes to the efficient dynamic alignment of cortical oscillatory activity to relevant phonological units in the speech signal (Giraud & Poeppel, 2012; Luo & Poeppel, 2007). Accordingly, greater language experience should be associated with more precise acoustic-temporal CTS (Lizarazu et al., 2021). Second, richer language experience should support the development of a neurocognitive model guiding the extraction of higher order linguistic information via lexico-semantic CTS (Broderick et al., 2021; ten Oever & Martin, 2021). Therefore, linguistic experience should also be positively associated with the cortical tracking of lexico-semantic information.

Language experience supports the development of language knowledge across linguistic domains (see also Gathercole & Thomas, 2005; Oller & Eilers, 2002; Paradis & Jia, 2016; Pérez-Navarro & Lallier, 2023). In particular, accumulated language experience enhances phonological acquisition (Nittrouer, 1996; Nittrouer & Burton, 2005; for a review, see Nittrouer, 2002) which, in turn, supports the comprehension of continuous speech. Of particular relevance for the purpose of our study, Goswami (2011, 2017) proposes that the cortical tracking of speech amplitude modulations within the speech envelope (acoustic-temporal CTS) supports phonological development during infancy and childhood. Within this framework, the accurate processing of the speech envelope is thought to guide and shape the emergence of phonological representations (Goswami & Leong, 2013; Leong & Goswami, 2014). Accordingly, a growing number of studies have now shown that acoustic-temporal CTS is at play as early as 4 months of age and developing throughout childhood (Attaheri et al., 2022; Menn, Ward, et al., 2022; Ríos-López et al., 2020).

Studies of atypical language development also suggest that CTS mediates the contribution of language experience to the acquisition of adequate linguistic abilities. In addition, several electrophysiological studies have shown that phonological deficits in children with dyslexia are tightly linked to atypical CTS within the delta and theta bands (Destoky et al., 2020; Di Liberto et al., 2018; Granados Barbero et al., 2022; Molinaro et al., 2016; Power et al., 2016) in comparison to both chronological-age-matched and reading-age-matched younger peers who are also matched on how much they have been exposed to written inputs (Di Liberto et al., 2018; Power et al., 2016). This suggests that impaired CTS could be causally implicated in atypical phonological development, and be somehow independent from linguistic experience (specially for written language). However, Destoky et al. (2020) reported poorer CTS in children with dyslexia only when compared to chronological and not to reading age-matched controls. Given these somewhat divergent findings, the extent to which accumulated language experience contributes to the maturation of CTS during phonological development remains unclear. Our study tackles this question by exploiting the largely different experiences that unbalanced bilingual children have within their two languages. Importantly, this group provides a unique opportunity to quantify the effect of accumulated language experience on acoustic envelope-level CTS and phonological development within the same participants. To gain a more comprehensive understanding of the relevance of CTS for the development of language skills, we go a step forward by exploring whether CTS is also linked to lexico-semantic knowledge through accumulated experience within a particular language.

Lexico-semantic knowledge, whose development is tightly related to accumulated language experience during childhood (Carbajal & Peperkamp, 2020; Oller et al., 2007; Paradis & Genesee, 1996; Pearson et al., 1997), has been linked to the efficiency of cortical oscillatory mechanisms for speech processing. Recent studies have shown that, in adults, lexico-semantic CTS modulated by both context-driven word predictability (Broderick et al., 2021; Klimovich-Gray et al., 2021; Koskinen et al., 2020; Molinaro et al., 2021). Of particular relevance for our hypotheses, these studies converge on the finding that the predictability of lexico-semantic information shapes CTS in adults, possibly as they have developed efficient neurocognitive language models as a function of accumulated language experience. However, the contribution of language experience to the development of lexico-semantic CTS during childhood is an open question that, to our knowledge, no study has directly addressed previously. By addressing such a question, we will be able to add evidence on a potential tradeoff between acoustic-temporal and lexico-semantic CTS from a developmental perspective.

Overall, the evidence reviewed above points to the relevance of acquired language knowledge through accumulated linguistic experience for the tuning of CTS at both the acoustic-temporal level (linked to phonology) and higher order linguistic levels (associated with lexico-semantics). Here, we study the relationship between input and CTS at both levels by capitalizing on a single group of unbalanced bilingual children, whose accumulated experience within their two languages varies greatly. Therefore, we are able to explore the role of greatly different language experiences on the cortical tracking of acoustic-temporal speech information (i.e., the envelope), and on the much less investigated mechanism of cortical tracking of lexico-semantic information in continuous speech. In addition, we assess whether the cortical tracking of these two types of speech information is related to phonological and lexico-semantic skills measured behaviorally.

## Method

### Participants

Participants were 35 (18 females) Basque-Spanish bilingual children between 6 and 7 years of age (*Mean age* = 6.92; *SD* = .11). We selected them from a broader longitudinal study (Pérez-Navarro & Lallier, under review) based on their amount of exposure to each language when they were 6.4 years old (the last stage of that longitudinal study). Namely, they were the group of children who had the most experience within their most exposed language (> 70% of their waking hours exposed to Basque, referred to as *Exp(+)* henceforth) and the least within their least exposed language (< 30% of their waking hours exposed to Spanish, *Exp(-)* henceforth). This selection criterion enabled us to investigate the influence of differential linguistic experience on CTS of two different languages within the same participants, thus limiting inter-subject variability for our language comparisons. Importantly, since participants had started formal reading instruction only a few months prior to participation in the study, the amount of written input was not sufficient to be considered a main source of language input at the time of testing. Thus, the potential influence of written language exposure on phonological abilities (Dehaene et al., 2015; Goswami & Bryant, 1990; Morais et al., 1979) and CTS in our study was considered to be minimal.

Participants had normal hearing, no history of neurological disorders, nor familial risk of developmental language disorder or any other cognitive-related genetic pathology. The study was approved by the BCBL Ethics Committee and complied with the Declaration of Helsinki.

### Behavioral session

The relative amount of linguistic experience within each language was assessed through a comprehensive linguistic questionnaire completed by the children’s parents. From this questionnaire, we extracted a composite index of the proportional amount of experience within each language (<u>Supplemental Formulas 1 to 3</u>). In addition to the amount of language experience, we assessed children’s phonological and lexico-semantic language knowledge.

Phonological abilities were evaluated with a nonword repetition task consisting of 18 items for each language, ranging from 3- to 6-syllable nonwords (6 nonwords per syllable length and language). In this task, participants were randomly presented with the 36 auditory nonwords, each constructed to follow either the stress and syllabic phonotactic constraints of Basque or those of Spanish. Upon participant’s oral repetition of each nonword, an experimenter coded each repetition as either correct or incorrect. Mean accuracy for each language was taken as a measure of language-specific phonological abilities.

Lexico-semantic knowledge was assessed through a picture-naming task consisting of 45 items for each language. The items were different in each language version, although matched in terms of word frequency. In each language version of the task, participants were asked to name the pictures that appeared on screen in a random order, without any time constraints. The experimenter coded each trial as either correct or incorrect upon participant’s response. The order of presentation of the two languages was counterbalanced across participants. In order to harmonize participants’ performance in the picture-naming task across both languages, we weighted performance on the different items (1, correct; 0, incorrect) by the inverse of their Zipf lexical frequency (van Heuven et al., 2014). For Basque words, lexical frequency was extracted from EHME database (Acha et al., 2014); and for Spanish words from EsPal database (Duchon et al., 2013). This way, more frequent (and presumably easier) words had a smaller weight on participants’ overall performance than less frequent ones that were less likely to be known by children. Thus, our measure of performance in the picture-naming task was Zipf-frequency-weighted accuracy.

Importantly, our phonological and lexico-semantic measures have previously proven sensitive to differential amounts of language exposure in bilinguals (Gámez et al., 2019; Paradis et al., 2016; Thordardottir et al., 2006).

### Electroencephalography (EEG) session

#### EEG task: speech listening

Participants listened to continuous streams of natural speech in the form of storytelling. We used two stories that were adaptations of two short books targeted at 6-year-old children, and followed a very similar narrative structure. Importantly, this literary style targeted at children offers a great number of different words and, consequently, considerable variability in their lexical frequencies (Nation et al., 2022; see Supplemental Figure 1), which was relevant to explore the effect of language experience and knowledge on the cortical tracking of lexico-semantic information.

The stories were about the history of outer space exploration and the evolution of life on Earth. For each story, both a Exp(+) (Basque) and a Exp(-) (Spanish) version was narrated by the same female native speaker of both languages in a child-directed speech register. Each story lasted about 15 minutes, because this duration has proven sufficient to robustly estimate cortical tracking of speech with EEG (Destoky et al., 2019). Half of the participants listened to the ‘outer space’ story in Basque, and the ‘life on Earth’ story in Spanish, and vice versa for the other half of participants. The order of language presentation was counterbalanced across participants. Participants were asked to listen attentively to the stories that were presented to them over speakers. They were sitting in a comfortable upright position and asked to look at static images depicting the story narrative, presented on the center of a computer screen positioned at 80 cm from their eyes. Every 5 minutes approximately, participants were asked three simple yes/no questions (9 per story in total) to check whether they were paying attention to, and comprehending, the stories.

#### EEG preprocessing

EEG data was recorded using a 64 Ag-AgCl electrodes standard setting (actiCAP, Brain Products GmbH, Germany). One electrode was placed over the outer canthus of each eye, and one below the left eye to monitor eye movements and blinks. Electrode impedance was always below 15 kΩ and remained below 10 kΩ in the vast majority of electrodes across participants. During data collection, raw EEG signal was amplified (BrainAmp DC, Brain Products GmbH, Germany), online high-pass filtered at 0.05 Hz, digitized using a sampling rate of 1000 Hz, and referenced to the midline central electrode (Cz).

To obtain the best possible temporal alignment between acoustic stimuli and EEG signal, we digitized the speakers’ acoustic signal, with a sampling rate of 1000 Hz (Polybox, Brain Products GmbH, Germany). This allowed us to compensate for varying lags between the digitized trigger and the actual presentation of acoustic stimuli. We then cross-correlated the amplitude values of the speakers’ acoustic signal and its corresponding audio template to ensure an optimal alignment. Thus, before EEG data preprocessing, triggers that marked the onset of each speech fragment were realigned to the time of maximum correlation with the actual presentation of the acoustic signal.

All EEG data preprocessing steps, and later data transformations were conducted at sensor level in MATLAB (version R2014B, MathWorks, 2014), using both custom code and functions from FieldTrip toolbox (version 20180604, Oostenveld et al., 2011). First, we downsampled EEG and audio signals to 200 Hz. Second, we bandpass filtered the signal between 0.2 and 40 Hz with a zero-phase fourth-order finite impulse response filter, using the default transition bandwidth in the FieldTrip toolbox for bandpass filtering. Third, we detected physiological artifacts through independent component analysis (ICA, runica method) on the filtered signal. After visual inspection of ICA, we subtracted independent components related to eye movements and blinks from the EEG signal (*mean number of rejected components per participant* = 2.06, *SD* = .5). Interpolation of bad channels was achieved using the weighted average of their neighbors. Our EEG data exclusion criteria was to not further analyze participant datasets for which more than 30% of the data was rejected. Although no participant exceeded such a threshold, two participants were excluded, as one of them did not want to remain seated while listening to the stories and the other fell asleep during the recording. Thus, we ended up with a sample of 33 analyzable EEG and behavioral datasets in each language.

For speech-brain coherence analyses, we divided the continuous EEG signal from each storytelling and language into 2000-ms epochs (the inverse of our lowest frequency of interest as well as frequency resolution, 0.5 Hz) with a temporal overlap of 1000 milliseconds. We discarded epochs and channels in which overall voltage departed more than 3 z-values from the average of all epochs and channels respectively (*mean percentage of epochs removed per participant* = 1.14 %, *SD* = .66; *mean number of channels removed per participant* = 1 out of 64, *SD* = 1.93). For mTRF analyses, we proceeded with the continuous EEG signal (without epoching).

#### CTS indexes

We used two types of CTS metrics (Figure 1). The first one was coherence (Halliday et al., 1995), which measures the phase correlation between two signals (here the brain and the speech signals, thus termed speech-brain coherence hereafter), which was used to extract the brain tracking of acoustic temporal speech information (i.e., the speech envelope). The second set of CTS metrics were extracted from the multivariate temporal response function (mTRF, Crosse et al., 2016) of continuous brain oscillatory activity to acoustic temporal and linguistic (i.e., lexico-semantic) information. mTRFs consist of the linear mapping of the values of continuous stimuli (acting as regressors) on continuous brain activity (acting as response, EEG in our case).

**Figure 1.**
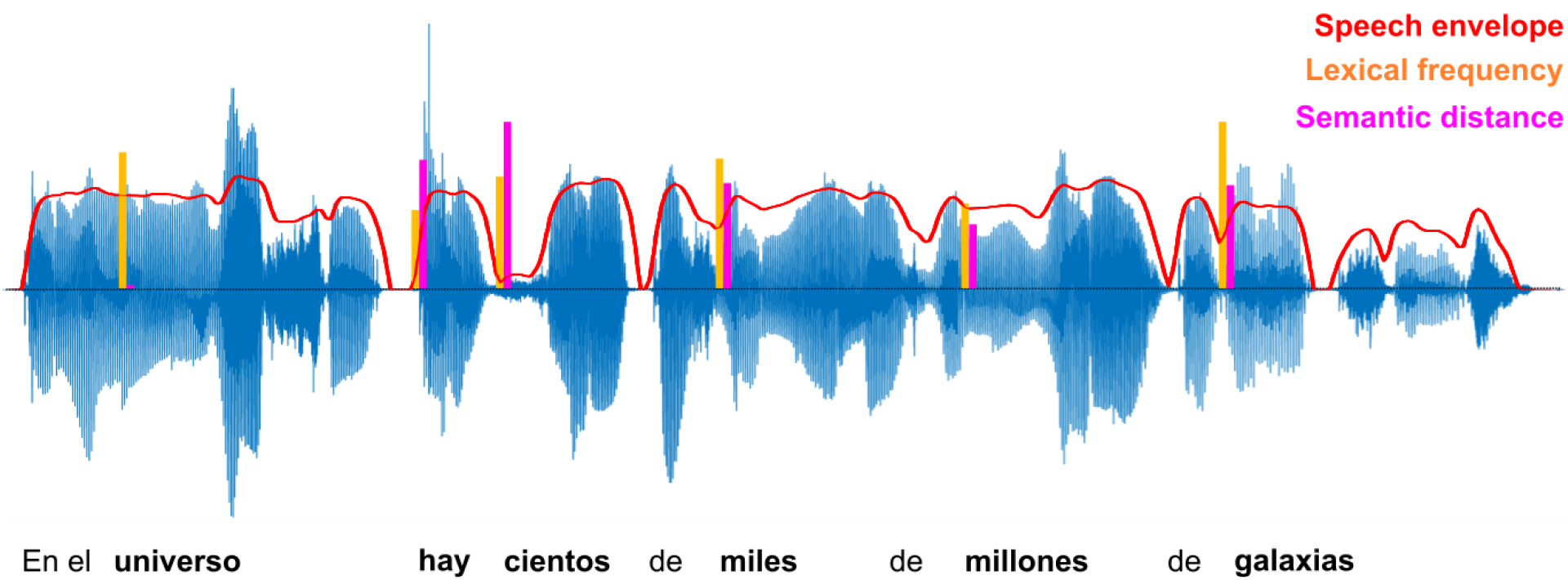
Graphical summary of the CTS analyses. In blue, the speech waveform of “In the universe, there are hundreds of billions of galaxies” in Spanish. Speech-brain coherence and envelope-level mTRF models are based on the relationship between the speech envelope (in red) and EEG activity. Lexical frequency and semantic distance mTRFs are obtained from the EEG response to bursts of different amplitude (lexical frequency, orange; sentence-level semantic distance, pink) at the onset of each content word.

##### Speech-brain coherence

We computed speech-brain coherence as the phase correlation between EEG and speech envelope (i.e., the Hilbert-transformed audio signal). To achieve this, we used custom functions in MATLAB following the specifications described in Molinaro and Lizarazu (2018). We circumscribed our speech-brain coherence analysis to the 0.5 – 10 Hz frequency range, which spans over the delta (0.5 – 4 Hz) and theta (4 – 7 Hz) frequency bands, which align to the timescales of prosodic phrasing (∼1000-2000 ms) and syllables (∼150-200 ms) respectively. In order to test whether speech-brain coherence within theta band was specifically related to syllable tracking, we estimated the syllable rate of the stories (based on an automatic algorithm, (de Jong & Wempe, 2009; in Praat, Boersma & Weenink, 2021). The overall average syllable rate was 5.63 Hz (*SD* = .33) and was highly similar in Basque (5.7 Hz, *SD* = .28) and Spanish (5.56 Hz, *SD* = .36). Given that our speech-brain coherence analysis had a frequency resolution of 0.5 Hz, the frequency bin that aligned most closely to the syllable rate was 5.5 Hz.

Coherence values vary between 0 (no linear phase relation) and 1 (total linear phase relation). To find the moment of maximum speech-brain synchronicity, we computed coherence between both signals at 6 different time lags (ranging from 40 to 140 ms in steps of 20 ms) in two arrays of sensors that have previously shown speech-brain coherence effects, located symmetrically within the left (i.e., T7, C3, TP7, and CP3) and right (i.e., T8, C4, TP8, CP4) temporal hemispheres. This a priori selection of sensors was only used to determine the time of maximum coherence, and not for localizing statistical coherence effects in our analyses (which were located through cluster-based permutation tests). A 60 ms lag of the EEG with respect to the speech signal was the time point of maximum coherence across participants and conditions, to which we circumscribed our speech-brain coherence estimates for statistical analyses.

##### mTRFs

We computed one mTRF encoding model per participant and language, which included four regressors: word onsets, the speech envelope, lexical frequency, and sentence-level semantic distance. Word onsets were employed as a control regressor, to account for the presumably high variability in the mTRF models associated with the mere responses to word onsets. Speech envelope served as a regressor for acoustic-temporal CTS, and lexical frequency and semantic distance for lexico-semantic CTS. Importantly, computing envelope-level mTRF also served to verify that it was related to the envelope tracking measured with coherence and to establish a positive control for mTRF analyses of lexico-semantic features, which are less salient in the signal. The speech envelope regressors have the same operationalization (i.e., the Hilbert-transformed audio signal) as in the speech-brain coherence analyses (also following the specifications in Molinaro & Lizarazu 2018).

Lexical and semantic regressors consisted of continuous vectors (one per language and story) containing bursts at the onset of every content word in each story. The amplitude of such bursts corresponded to latent variables that were used as proxies for lexical and semantic information. For the lexical regressor, we used lexical frequency: the amplitude of each burst was the inverse of the Zipf lexical frequency of its corresponding word (van Heuven et al., 2014; see Supplemental Figure 1). For the semantic regressor, we computed sentence-level semantic distance vectors following a similar approach to Broderick et al. (2021; see Supplemental Figure 2). We first obtained the semantic representation of each story word in a 300-dimensional space through the fasttext Python package for text representations (Joulin et al., 2016). Then, the amplitude of each burst was computed as 1 minus the Pearson correlation of the semantic dimensions of a word with the average of all its preceding words within a sentence. This way, lexical items with a bigger semantic correlation with the preceding words within a sentence had lower amplitude bursts, and less semantically related words had higher amplitude bursts relative to their preceding context.

After obtaining all the regressors of interest, we used the mTRF toolbox (Crosse et al., 2016) in MATLAB to fit one mTRF encoding model with the EEG response with all features (envelope, lexical frequency, and sentence-level semantic distance), and for each language and participant. Our mTRF analysis time window was 900 ms, spanning from -150 ms pre- to 750 ms post-stimulus. In order to train and test the encoding model, we split our continuous EEG signal and each corresponding feature vector (∼ 16 minutes) into 8 folds of equal length (∼ 2 minutes). We trained the encoding model in 7 folds and tested its accuracy on the remaining one, repeating this process for 30 iterations per fold. We then extracted two mTRF metrics for between-languages comparisons of CTS. First, we assessed whether and to what extent each individual regressor contributed significantly to the model fit (through *prediction correlation*, r-values). Thus, within 30 iterations per fold for each regressor of interest (envelope, lexical frequency, and semantic distance), we contrasted the full model fit (all regressors) versus the fit of the model where one regressor was permuted, and averaged such r-values across iterations. The resulting fold-average r-values, the above-chance prediction correlation coefficient between each feature, and the mTRF (at electrode level) were used in further statistical analyses as proxies for the extent to which each feature was linear mapped by EEG activity and to confirm that this feature contributed to the model-fit above the chance level across all subjects (see mTRF group analysis section). For the main analysis, the prediction correlation coefficient (the difference between the full model r-value and the specific feature-permuted model r-value) served as an electrode-level estimate of how faithfully EEG activity mapped a given speech feature (e.g., lexical information).

The second mTRF metric was the *temporal weights* of the EEG response to each regressor. Such weights form an ERP-like response that indexes EEG temporal sensitivity to changes in a given regressor (see Crosse et al., 2021). Thus, the temporal weights allowed us to test whether there were relevant between-languages differences in the amplitude or the latency of the temporal response to the envelope, lexical, or semantic information.

#### Statistical analysis

##### Behavioral language measures

In order to assess between-languages differences in language performance, we fitted linear mixed effect (LME) models with language (Exp(+), Exp(-)) as predictor for each behavioral language measure (language experience, vocabulary knowledge, and phonological ability), and participants as random intercepts, to account for between-participant differences. LME models were fitted using the lmer function from lme4 package (Bates et al., 2015) in R (version 4.2.1, R Core Team, 2022). In addition, we used Bayesian t-tests in JASP (version 0.16.4, JASP team, 2022) to test the strength of the evidence for between-languages similarities in our behavioral measures in the case of non-significant between-languages differences.

##### Speech-brain coherence

We used cluster-based permutation tests (CBPTs) to statistically estimate speech-brain coherence. CBPT is an efficient way of estimating the presence of a statistical effect in a high dimensional space, as it is able to account for the spatial adjacency of electrodes and test for significant effects that are shared across a group of electrodes (a cluster) (Maris & Oostenveld, 2007). In our case, we ran dependent-samples two-tailed CBPTs in FieldTrip with 1000 permutations. Our CBPTs required at least 2 electrodes to form a cluster as a way to limit the possibility of single-electrode false-alarm effects, and we corrected for multiple comparisons based on the number of a-priori significant clusters. We first assessed whether there was above-chance speech-brain coherence in each language by contrasting through CBPT the phase alignment of EEG and genuine speech envelope versus the phase alignment of EEG and a surrogate version of the envelope that did not follow the original speech order (i.e., flipped speech envelope surrogate). Then, we contrasted through CBPT the speech-brain coherence values of Exp(+) and Exp(-) to analyze whether there was a significant between-languages effect on this CTS metric.

##### mTRFs

Before our analyses of interest, we conducted a control test to verify that, at the group level, there were significant prediction correlations between our three regressors of interest and the mTRF model. Namely, we conducted a one-sample t-test (one-tailed) against the null hypothesis (no different prediction correlation from 0) for each of the regressors. After such control, and similarly to when contrasting speech-brain coherence between languages, we compared Exp(+) and Exp(-) through 6 CBPTs, one per mTRF metric (i.e., prediction correlation and temporal weights) and regressor (i.e., envelope, lexical frequency, and semantic distance). Both prediction correlation and temporal weights were analyzed at electrode level.

##### Relationship between CTS and behavioral language measures

In order to assess the correspondence between CTS and behavioral language performance in the phonological and lexico-semantic domains, we fitted two LME models, one with acoustic-temporal CTS and one with lexico-semantic CTS as outcome measure. In the first LME model, phonological abilities and vocabulary knowledge, as well as their interaction with language, served as predictors of acoustic-temporal CTS, a composite of speech-brain coherence and envelope-level TRFs. In the second LME model, the same set of measures were employed as predictors of lexico-semantic CTS: a composite score of lexical- and semantic-level TRFs. We included participants as random intercepts in both LME models, which allowed us to account for baseline individual differences in the different languages and language domains. As for our behavioral language analyses, we used the lmer function from lme4 package in R. In order to test for omnibus main effects and interactions of the predictors, we used the anova function in base R.

## Results

In line with their significantly higher language experience (i.e., percentage of exposure) within Exp(+) than within Exp(-), *t*(63) = 35.13, *p* < .001 (*β* = 72.298, *SE* = 2.058), participants showed significantly higher vocabulary knowledge in Basque than in Spanish, *t*(32) = 13.18, *p* < .001 (*β* = 0.077, *SE* = .006; see Figure 2). However, there was no between-language difference in phonological abilities, *t*(30.1) = -0.84, p > .05 (*β* = -0.014, *SE* = .017), nor in the comprehension of the stories that participants listened to during the EEG session, *t*(28) = 0.12, *p* > .05, (*β* = 0.005, *SE* = .039). Indeed, Bayesian t-tests yielded moderate evidence for between-language similarities in phonological abilities, *BF_10_* = .285, *error* = .034 %, *median difference* = -0.151, *CI* [-0.492 .184], and story comprehension respectively, *BF_10_* = .22, *error* = .024 %, *median difference* = -0.04, *CI* [-0.417 .334].

**Figure 2.**
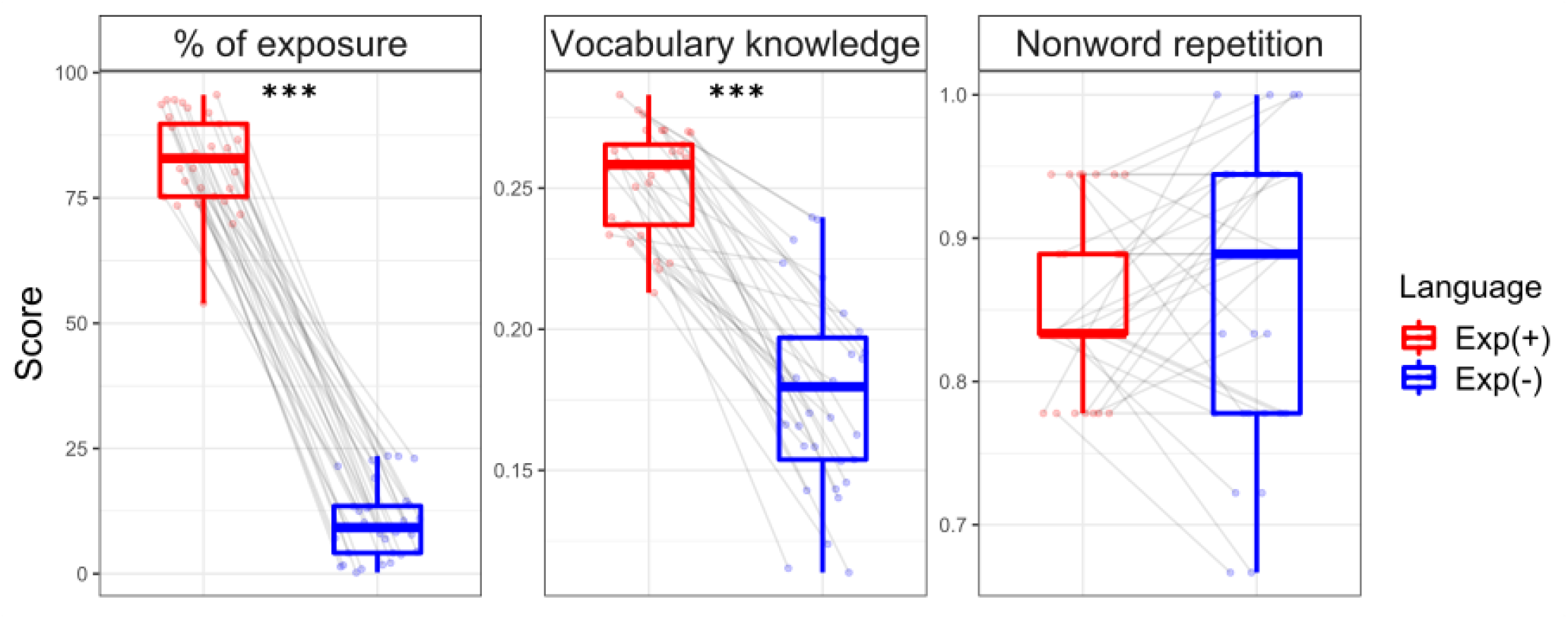
Amount of linguistic exposure and performance on vocabulary (picture naming) and phonology (nonword repetition) tasks in each language. Points represent each individual score in the different measures and languages. Boxplots represent group estimates, with horizontal lines within each box marking the median score. Upper and lower hinges mark the first and third quartile, and whiskers show 1.5 * interquartile range. Lines connect the scores of each participant across both languages. Asterisks indicate a significant difference between languages (***p < 0.001).

### Decreased language experience boosts acoustic-temporal CTS

#### Coherence

Within the delta frequency band, speech-brain coherence was significant between 0.5 and 1.5 Hz in Exp(+), *cluster statistic* = 271.73, *p* < .001, *SD* = .001, and Exp(-), *cluster statistic* = 278.17, *p* = .001, *SD* = .001 (Figure 3, A). However, there were no between-language difference. Coherence in this delta range had considerably overlapping topographies in both languages (Figure 3, B). In the theta range (4 – 7 Hz), we did not find significant speech-brain coherence in any of the languages (all cluster-corrected *p-values* > .05). We also did not observe significant coherence in the specific 5.5 Hz bin that aligned closely to the syllable rate of both languages (*p* > .05).

**Figure 3.**
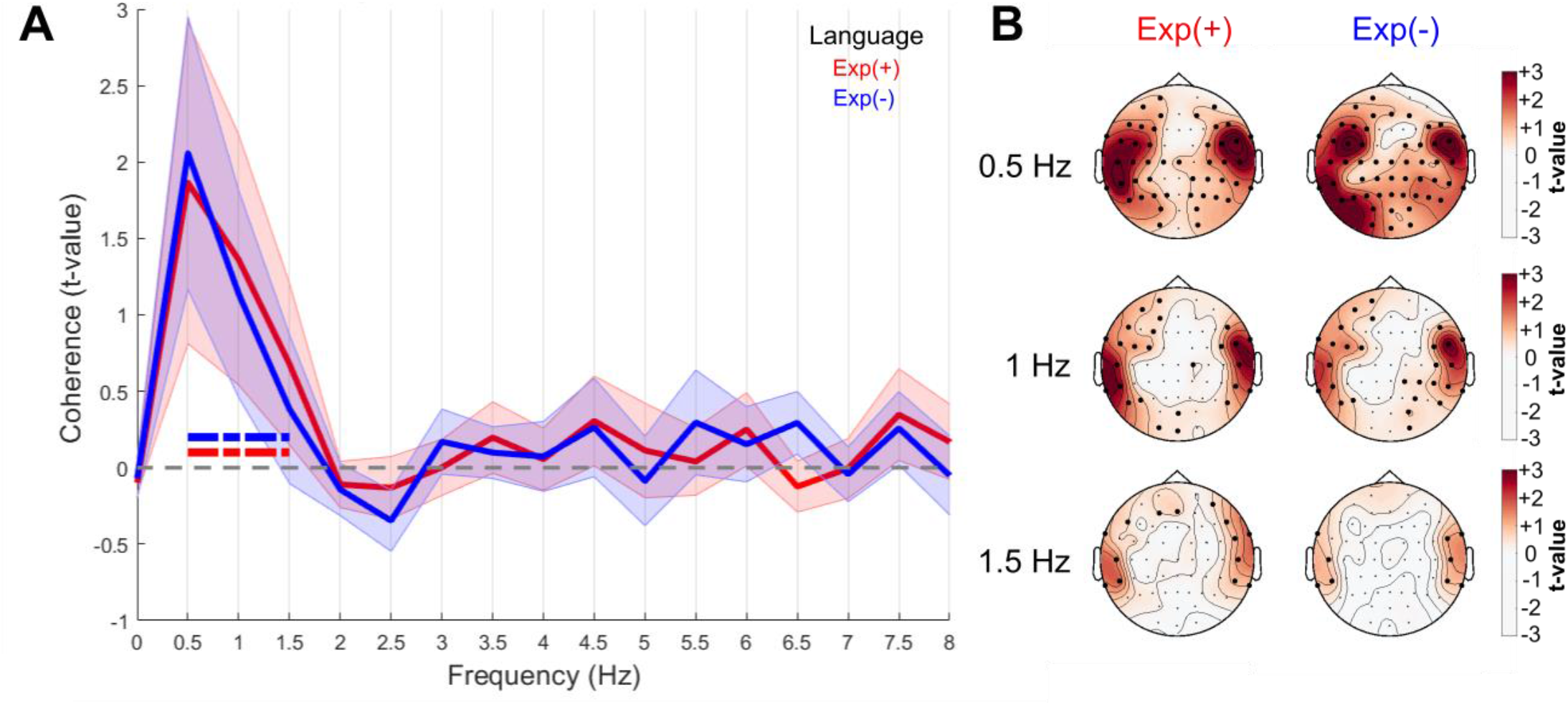
A) Speech-brain coherence across frequency bands for Exp(+) (red), Exp(-) (blue) against their random surrogate (flipped version). The discontinuous horizontal lines on the bottom mark the frequency range in which there was significant coherence for Exp(+) (red), and Exp(-) (blue). Red and blue shaded areas represent the standard error. B) Topography of speech-brain coherence in the 3 significant frequencies in Exp(+) (left) and Exp(-) (right). The colormap marks the size of the difference in coherence (normalized) between genuine speech and its flipped version (i.e., red, higher coherence; white, lower). Bigger black circles within each topographic map signal electrodes with significant speech-brain coherence from CBPT.

#### mTRF

Our control tests, one-sample t-tests (one-tailed) against no different prediction correlation from 0, showed that our three regressors of interest contributed above chance level to the mTRF model (speech envelope: *t*(65) = 12.78, *p* < .001, *d* = 1.57; lexical frequency: *t*(65) = 6.50, *p* < .001, *d* = .8; semantic distance: *t*(65) = 5.27, *p* < .001, *d* = .65; Supplemental Figure 3), which enabled us to conduct between-languages comparisons of their fit.

Regarding acoustic-temporal CTS, CBPT yielded significantly higher prediction correlation coefficients (Pearson’s r) between envelope and EEG mTRF in the less exposed language (Exp(-)) than in the more exposed one (Exp(+)), *cluster statistic* = -27.65, *p* = .01, *SD* = .004 (Figure 4, B). Regarding the temporal weights in response to the speech envelope, CBPT did not show significant differences between languages (*cluster-corrected p-values* > .05; Figure 4, A). This suggests differences in how strongly children rely on envelope tracking between their two languages (i.e., stronger reliance on envelope tracking in their less experienced language), but no between-language differences in the temporal characteristics of the response to the envelope.

**Figure 4.**
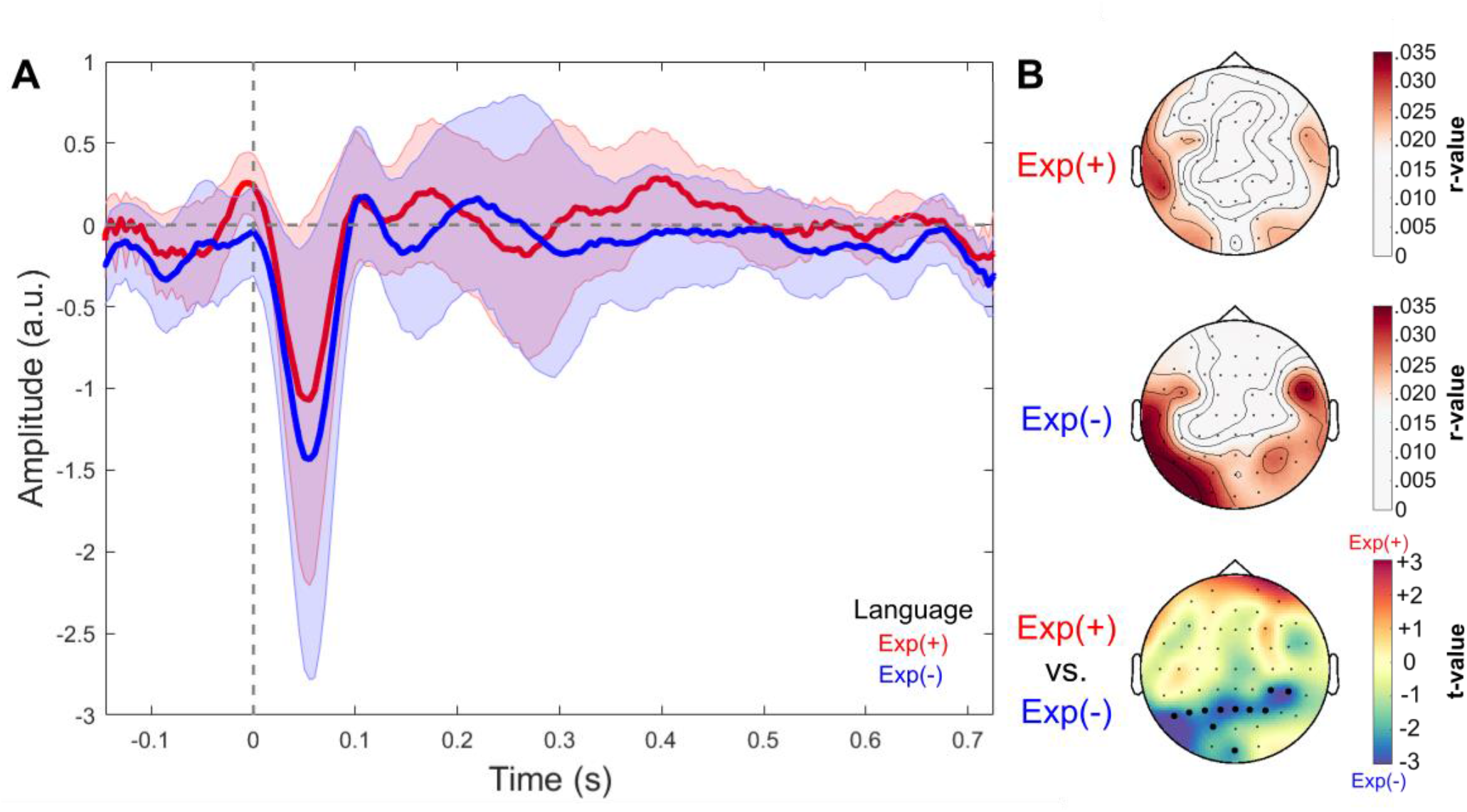
A) Temporal weights of the EEG signal TRF to the speech envelope in Exp(+) (red) and Exp(-) (blue). Red and blue shaded areas represent the standard error. B) Topography of the prediction correlation coefficients (r-values) between speech envelope and EEG TRF for Exp(+) (top), Exp(-) (middle), and their difference (bottom). The colormap for the two top topographies marks the degree of correlation (r-values); while the bottom topography is color mapped according to the size of the difference (t-value) between Exp(+) (relatively higher correlation coefficients in red) and Exp(-) (relatively higher correlation coefficients in blue). Bigger black circles signal electrodes in which CBPT yielded a significantly bigger prediction correlation at envelope level in Exp(-).

#### Language experience increases semantic sensitivity by CTS

CBPT did not yield significant between-language differences in the prediction correlation coefficients (model-fit r values derived from mTRF) between lexical frequency and EEG activity (*cluster-corrected p-values* > .05). The temporal weights in response to lexical frequency showed a similar latency to lexical-level TRFs in adults (e.g., Broderick et al., 2021) and did not show significant between-language differences (*cluster-corrected p-values* > .05; Figure 5, A).

**Figure 5.**
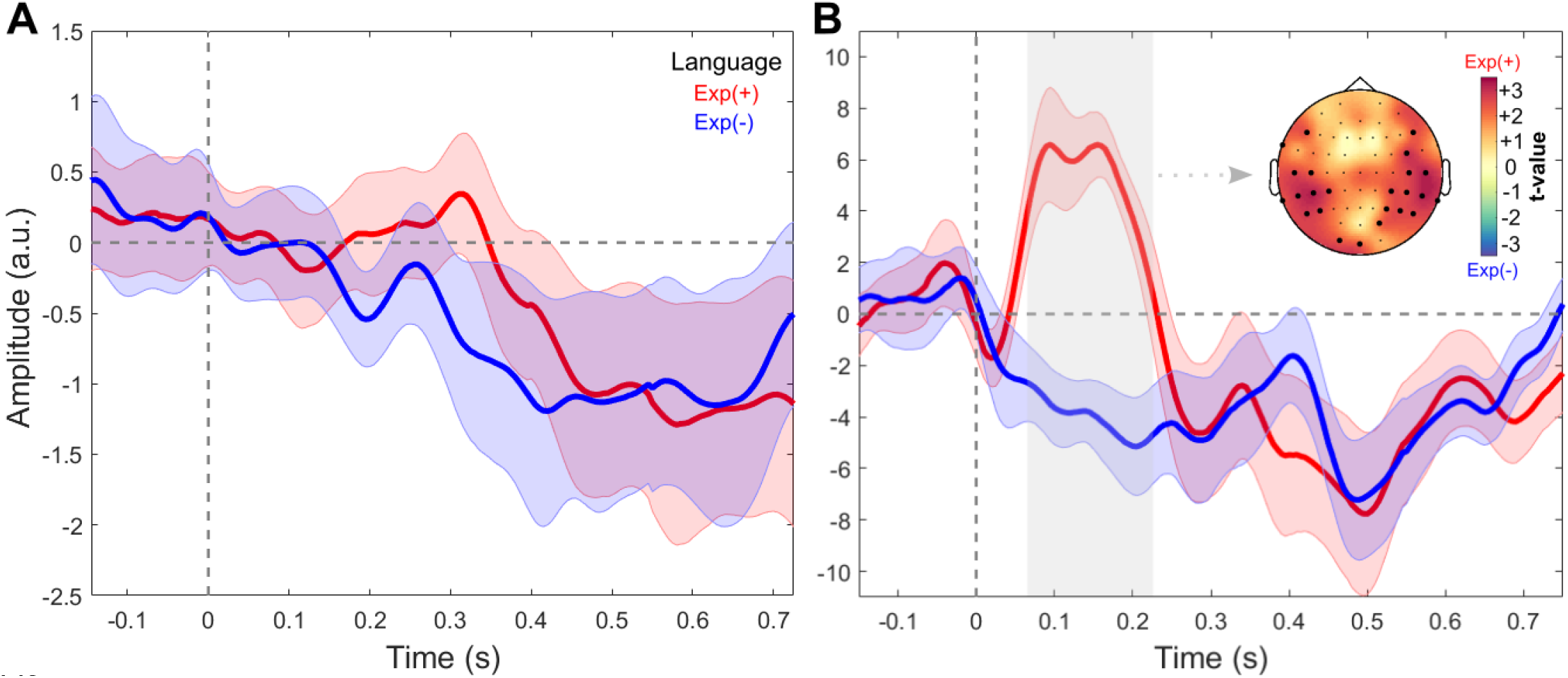
A) Temporal weights of the EEG signal TRF to lexical frequency in Exp(+) (red) and Exp(-) (blue). Red and blue shaded areas represent the standard deviation of the temporal weights. B) Temporal weights of the EEG signal TRF to semantic distance in Exp(+) (red) and Exp(-) (blue). Red and blue shaded areas represent the standard error of the temporal weights. The gray rectangle marks the latency of significant between-languages differences. The distribution over the scalp of such a significant difference in semantic distance (Exp(+) > Exp(-)) in this time window is represented by bigger dots in the top right topographic map.

Regarding EEG responses to semantic distance, we found relevant differences between languages. While mTRF prediction correlation coefficients (r-values) for semantic distance did not differ between languages (*cluster-corrected p-values* > .05); CBPT yielded a significant difference in the temporal response (i.e., weights) to semantic distance between the Exp(+) and Exp(-) languages (*cluster statistic* = 2827.4, *p* = .009, *SD* = .003). Thus, within this group of participants, EEG signal showed significantly higher sensitivity to semantic information in Exp(+) than in Exp(-) between 70 and 230 milliseconds after word onset (Figure 5, B). This suggests earlier tracking of the relevant contextual-semantic information for the language to which bilingual children were exposed the most.

#### Acoustic-temporal CTS is specifically linked to phonological abilities, lexico-semantic CTS to vocabulary knowledge

Acoustic-temporal CTS and lexico-semantic CTS composite scores were computed by aggregating EEG metrics for a more robust estimate of CTS-behavior relationships, while limiting the possibility of inflating type I error due to multiple tests. More specifically, an acoustic-temporal CTS composite was composed by averaging the normalized values of: 1) speech-brain coherence at delta (0.5-1.5 Hz), 2) speech-brain coherence at theta (4-7 Hz), and 3) prediction correlation values for envelope TRF. The lexico-semantic CTS composite consisted of: 1) prediction correlation values for lexical frequency TRF, and 2) prediction correlation values for semantic distance TRF. These composite scores were validated by the strong positive correlations found between (i) coherence to the speech envelope at delta and theta, and the envelope mTRF (*average Pearson’s r coefficient* = .59; Exp(+): .6; Exp(-): .57; all *p-values* < .01) and (ii) lexical frequency and semantic distance mTRFs (*r* = .79; Exp(+): .81; Exp(-): .78; all *p-values* < .01).

Regarding the relationship between acoustic-temporal CTS and behavioral language measures, there was a significant interaction between phonological abilities and language as predictors of acoustic-temporal CTS [*F*(2, 46.52) = 4.69, *p* < .014]. This interaction was driven by the significant and positive relationship between acoustic-temporal CTS and phonological abilities (nonword repetition) in Exp(+) [*t*(45.69) = 3.05, *p* = .004 (*β* = .62, *SE* = .21)], but not in Exp(-) [*t*(48.93) = .62, *p* > .05 (*β* = .09, *SE* = .14)] (Figure 6). There was no significant relationship between envelope-CTS and vocabulary knowledge in either of the two languages (all *p-values* > .05).

**Figure 6.**
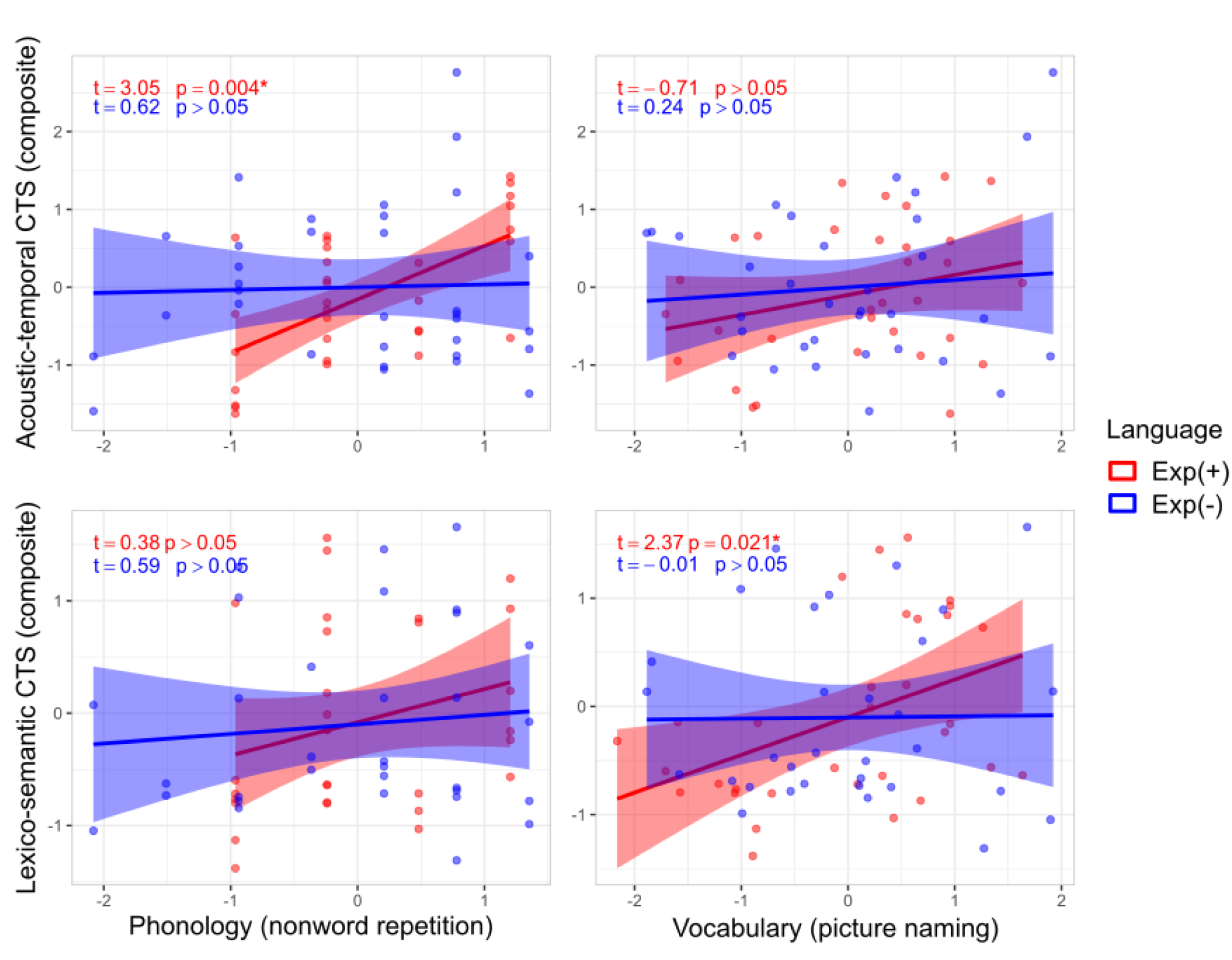
Relationship between CTS (acoustic-temporal and lexico-semantic) and language knowledge (phonology and vocabulary). Lines represent the slopes of linear regressions between CTS and behavioral metrics in each language respectively, with shaded areas around the lines marking 95% confidence intervals. Estimates in top left corners are the t- and p-values of linear mixed effect models for each relationship; asterisks indicate a significant effect.

In parallel to the relationship between phonological abilities and envelope-level CTS, we found that vocabulary knowledge was positively related to lexico-semantic CTS. In this case the interaction between vocabulary knowledge and language was only marginally significant [*F*(2, 57) = 2.81, *p* = .068]. Nonetheless, this interaction was also driven by an underlying significant and positive relationship between vocabulary knowledge and lexico-semantic CTS only in Exp(+), *t*(57) = 2.37, *p* = .021 (*β* = .41, *SE* = .17), and not in Exp(-), *t*(57) = -0.006, *p* > .05 (*β* = -0.001, *SE* = .14) (Figure 6). There was no significant relationship between lexico-semantic CTS and phonological abilities in either of the two languages (all *p-values* > .05).

## Discussion

The main goal of the present study was to estimate the impact of early linguistic experience on the cortical tracking of speech, by assessing the CTS of 6-year-old unbalanced bilingual children in both their languages. We found relevant differences between the more experienced language (Basque) and the less experienced language (Spanish) in the cortical tracking of acoustic-temporal (speech envelope) and lexico-semantic speech features. Such differences can be understood from a developmental perspective in which linguistic input plays a relevant role in the maturation of the cortical tracking of different types of linguistic information encoded in the speech signal. Additionally, and given that CTS was linked to language knowledge (in vocabulary and phonology) at behavioral level, our results inform the relevance of CTS for indexing the development of specific language abilities. Moreover, the present study is, to our knowledge, the first to assess the cortical tracking of two different languages (and at different levels of acoustic-linguistic complexity) within the same group of bilingual children.

We found robust acoustic-temporal CTS in both of the languages of bilinguals through two different metrics, namely speech-brain coherence and envelope-level mTRF models. Significant speech-brain coherence was found within the 0.5 to 1.5 Hz delta range, the timescale of prosodic stress occurrences, whose tracking has been shown to be relevant for efficient lexical segmentation (Kooijman et al., 2009). In line with previous research in children (Ríos-López et al., 2020), we did not find significant theta-band speech-brain coherence, which has been related to the tracking of syllabic speech units in continuous speech and might be more relevant at later stages of development, such as adulthood (Doelling et al., 2014; Peelle et al., 2013). It is possible that the developing language system of children is more sensitive to the slower delta frequency timescales for tracking continuous speech, and only starts exploiting faster syllabic amplitude modulations once their phonological representations are more developed, possibly through sufficient experience and proficiency. Indeed, a recent cross-linguistic study by Peter et al. (2022) showed that acquired language-specific knowledge influenced the cortical tracking of syllabic information in adults. They demonstrated that adult native speakers of French (a syllable-timed language) showed enhanced cortical tracking of the syllable rate (∼5 Hz) compared with English and Japanese listeners (stress- and mora-time languages respectively), for whom syllable-timing might be less relevant for extracting phonological information.

However, in another recent study, Menn et al. (2022) did find significant speech-brain coherence at the syllable rate in infants as young as 9 months of age. Task requirement differences could explain these divergent findings between Menn et al. (2022) and our study at the theta frequency band. In Menn and collaborators’ study, infants’ CTS was measured while they were listening to their mothers in relatively short (40 second) live interactions (audiovisual stimuli), whereas in our study, CTS metrics were extracted from considerably longer 15-minute continuous stories with static images (very limited visual information). It is therefore possible that the length of the tasks and the use (or lack) of additional visual cues modulated CTS differently between studies, in line with the attested supportive effect of visual information for theta-band cortical tracking of speech (Lakatos et al., 2008; Power et al., 2013).

The lack of significant between-language differences in speech-brain coherence analyses shows that the young children in our study robustly tracked the speech envelope in both their more and less experienced languages. Nonetheless, when it comes to envelope-level TRFs, we found that linguistic experience did shape children’s cortical tracking of speech envelope information. Namely, the continuous EEG activity of this group of children encoded the speech envelope of their less experienced language (Spanish) more faithfully than that of their more experienced one (Basque). This was observed despite the fact that, at the time of testing, children had developed similar receptive language abilities in both their languages, illustrated by the lack of relevant between-language differences in phonological abilities and in story comprehension.

These findings help contextualize the role of cortical tracking of the speech envelope for language development. During language acquisition, envelope tracking is tightly linked to phonological development and broader language outcomes (Goswami et al., 2010; Kalashnikova et al., 2019; Vanvooren et al., 2017). Relatedly, envelope CTS by adult second-language learners increases with second language proficiency (Lizarazu et al., 2021). Therefore, the fact that the children encoded envelope-level information more faithfully in their less experienced language suggests they relied to a larger extent on low-level acoustic information to achieve a similar level of comprehension as in their more exposed language. We propose that acoustic-temporal CTS could thus serve as a route for speech comprehension during language learning, possibly by compensating while increasing higher-order linguistic knowledge is being learned. Thus, it is likely that as individuals can exploit an increasingly proficient language model for CTS, they gradually shift from relying on low-level temporal information towards tracking higher order linguistic units and structures.

Beyond the speech envelope, we report initial evidence on the cortical tracking of abstract linguistic features from continuous speech in children. Such evidence provides a developmental perspective to the role of language knowledge in the cortical tracking of higher order linguistic information, which is being increasingly studied in adults (e.g., Broderick et al., 2021; Kaufeld et al., 2020; Reetzke et al., 2021). Specifically, we found that children showed an early increased temporal sensitivity to semantic-level information only in their more experienced language (Basque). In addition to an expected peak between 400 and 600 ms that did not differ significantly between languages (Broderick et al., 2021), the temporal response to semantic distance showed a peak between 70 and 230 ms in the more experienced language (Basque), but not in the less experienced one (Spanish). We did not expect the cortical response to the semantic context to show a peak at such early latency. However, it falls within a similar latency range (∼50-200 ms) to several adult studies that found early contextual (typically semantic) influence on the cortical tracking of continuous speech (Brodbeck et al., 2018, 2023; Broderick et al., 2019; Donhauser & Baillet, 2020).

A plausible interpretation of such early sensitivity to semantic information is that, when children have more established language knowledge (only in the more experienced language here), they can capitalize on the semantic context to predict and efficiently track the phonological and/or lexical information from upcoming speech. Two interrelated factors suggest that, in the case of our study, such difference between languages in the cortical exploitation of semantic information might be driven by lexical, but not phonological, facilitation. First, Basque and Spanish overlap to a great extent in their phonological systems, but not in their vocabularies (Ezeizabarrena & Alegria, 2015). Second, the children showed similarly developed phonological abilities in both languages, but significantly greater lexical knowledge in their more experienced language. Therefore, it is plausible that their increased cortical sensitivity to contextual semantic information in their more experienced language is related to a more efficient lexical (rather than phonological) processing of the upcoming word in continuous speech. This interpretation is consistent with the positive relationship found between the cortical tracking of lexico-semantic information and vocabulary knowledge.

The relevance of acoustic-temporal and lexico-semantic CTS for explaining behavior was evaluated by assessing their link to individual differences in language knowledge within both phonology and vocabulary domains. Indeed, there was a specific relationship between acoustic-temporal (envelope) CTS and phonological abilities, only in the more experienced language (Basque). This finding is in line with Molinaro et al., (2016) and Di Liberto et al. (2018), who respectively reported that the cortical tracking of the envelope and phonetic features were related to phonological abilities in dyslexic children. Moreover, our finding is coherent with the hypothesis that sensitivity to prosodic and syllabic speech AMs (assessed directly here through speech-brain coherence) subserves the temporal sampling of phonological units within these timescales (phrases, words, and syllables) during phonological development (Goswami, 2011, 2017; Goswami & Leong, 2013; Leong & Goswami, 2014). Accordingly, bilingual participants who exhibited higher CTS to such speech AMs were those who showed more developed phonological processing abilities. Thus, our results support that phonological abilities and acoustic-temporal CTS develop hand in hand (Boets et al., 2007; Di Liberto et al., 2018; Lundberg et al., 1988; Power et al., 2016; Vanvooren et al., 2017), with CTS also predicting later reading outcomes (Ríos-López et al., 2021).

It is relevant to note that, while tracking speech AMs contributes to the development of phonological skills (Leong & Goswami, 2014, 2017; Molinaro et al., 2016), studies in adults highlight that CTS detaches from faithfully tracking the envelope, at least in normal and effortless listening environments (Hauswald et al., 2022; Schmidt et al., 2023). Therefore, the role of acoustic-temporal CTS for language development might decrease as soon as individuals can rely on sufficient lexico-semantic knowledge to track continuous speech (Kaufeld et al., 2020; Molinaro et al., 2021). A complementary possibility is that, along with language development, delta oscillatory activity shifts gradually from acoustic-temporal CTS (at envelope-level) towards semantic and syntactic CTS. This possibility aligns with a suggested developmental shift when exploiting prosodic information, namely from recognizing word boundaries early in life towards extracting semantic and syntactic information later in childhood (Friederici & Männel, 2014; Guasti, 2002; Männel et al., 2013). Indeed, Kaufeld et al. (2020) found that linguistic content modulated delta neural oscillatory activity beyond stimulus-driven prosodic timing. It is therefore possible that, during language development, delta CTS helps map the acoustically driven prosodic timing during language learning, while concurrently allowing the accumulation of knowledge about the linguistic units and structures present at similar acoustic timescales, such as syntactic information. Interestingly, Menn, Ward, et al. (2022) reported that 10-month-old infants delta (prosodic) CTS predicted vocabulary knowledge at 24 months. It is therefore possible that, in our study, children relied more on extracting envelope-level phonological information in their less familiar language, hence the significantly stronger prediction correlation of envelope-level TRFs in that language; and utilized the same delta tracking to extract increasingly more abstract linguistic information in their more exposed language. Future longitudinal studies assessing the different reliance on acoustic-temporal or linguistic features for tracking the speech signal as a function of age and language knowledge, will help clarify whether and when this potential developmental shift takes place.

Parallel to the acoustic-temporal CTS-phonology relationship, we observed a significant positive relationship between lexico-semantic CTS and vocabulary knowledge, only in the most experienced language (Basque). Indeed, these results provide initial evidence for the behavioral relevance of the cortical tracking of lexico-semantic information during language development. Interestingly, CTS was linked to both phonological and vocabulary performance only in the more experienced language, suggesting that the more stable language representations and processes in this language might map more consistently and faithfully onto individual differences in CTS. In contrast, weaker knowledge in the less experienced language might not involve as efficiently the neural resources for CTS. Accordingly, the language knowledge of children was overall more variable in their less experienced language (Figure 2). It is also plausible that significant behavioral-CTS relationships in the less exposed language had been detected with a larger sample size, by accounting for the higher intra- and inter-individual variability.

Overall, the present study informs theoretical, empirical, and computational accounts of the relevant role that linguistic experience plays in continuously shaping cortical oscillatory activity mechanisms that extract linguistic information from speech. Given our and previous findings, we propose that acoustic-temporal (envelope) CTS serves as phonological route for speech processing while a language is being learnt. Later on, and as a function of maturational factors such as age and linguistic experience, expert neurocognitive language models start exploiting lexical, semantic, and syntactic information to organize linguistic meaning (Martin, 2020; Martin & Doumas, 2017), group words within or between phrases (Meyer et al., 2017), and predict upcoming linguistic information (Molinaro et al., 2021; ten Oever & Martin, 2021).

In summary, our findings shed new light on the role that linguistic experience plays in the cortical tracking of speech, a neurocognitive mechanism supporting speech comprehension. For the first time, we show that, during language development, the maturation of the cortical tracking of the temporal speech envelope and lexico-semantic speech features is tightly linked to the amount of exposure to a language and the subsequent knowledge attained in that language. We also highlight the behavioral relevance of CTS, by showing that individual differences in specific language abilities are linked to the cortical tracking of specifically related linguistic features in continuous speech.

## Limitations of the study

There are two main study limitations that could be helped by future research. First, our single timepoint study only allows us to observe that acoustic-temporal and lexico-semantic CTS is indeed sensitive to differences in language exposure. However, longitudinal data would help us further understand whether the developmental tradeoff that we point towards is in place. In other words, whether the shift from relying on acoustic-temporal towards lexico-semantic information for tracking continuous speech takes place as a function of accumulated linguistic exposure in addition to other crucial maturational factors for language development, such as chronological age.

Also, a bigger sample size would have allowed us to compensate for the higher variability in behavioral language knowledge in the less experienced language (Figure 2), and test whether the specific links between CTS and behavioral language knowledge take place also in the second languag. A minor caveat of our study, specific to the topographic interpretation of our results, is that our reference electrode was located on the scalp midline central position (Cz), which could have boosted the signal-to-noise ratio for sensors located apart from such position. And, thus, yield more observable effects towards lateral temporal and parietal areas.

## Acknowledgements

Data, code, and materials necessary to reproduce the analyses presented here will be made publicly accessible at the Open Science Framework <u>repository</u> of the project upon publication. The analyses presented here were not preregistered. We would like to thank the participants and their families, as well as the BCBL lab department, for their contribution and commitment over the years of this longitudinal study. Special thanks to Araitz Garnika, Leire Eizagirre, Ioanna Taouki, for their support with stimuli preparation and EEG data collection, to Giovanni Di Liberto for his support with mTRF modeling, and to Caroline Handley for her proofreading and suggestions on this manuscript. This research was supported by the FPI grant BES-2016-078125 (to ML and JPN) and by the PID2022-136989OB-I00 (to ML) by Ministerio Español de Economía, Industria y Competitividad MINECO and Fondo Social Europeo; through project RTI2018-096242-B-I00(MCIU/AEI/FEDER, UE) funded by Ministerio de Ciencia, Innovación y Universidades (MCIU), the Agencia Estatal de Investigación (AEI) and Fondo Europeo de Desarrollo Regional (FEDER); by the Basque Government through the BERC 2018-2021 program and by the Spanish State Research Agency through BCBL Severo Ochoa excellence accreditation SEV-2015-0490. The authors declare no competing interests.

## CRediT authorship contribution statement

**Jose Pérez-Navarro**: Conceptualization, Methodology, Formal analysis, Writing – Original draft, Writing – Review & Editing Investigation, Data curation, Visualization, Funding acquisition; **Anastasia Klimovich-Gray**: Methodology, Formal analysis, Writing – Review & Editing, Data Curation, Resources, Visualization; **Mikel Lizarazu**: Methodology, Formal analysis, Writing – Review & Editing, Data Curation, Resources, Visualization; **Giorgio Piazza**: Data collection, Data curation, Writing – Review & Editing; **Nicola Molinaro**: Conceptualization, Writing – Original draft, Writing – Review & Editing, Supervision; **Marie Lallier**: Conceptualization, Writing – Original draft, Writing – Review & Editing, Supervision.

## Supplemental material

**Supplemental Figure 1.**
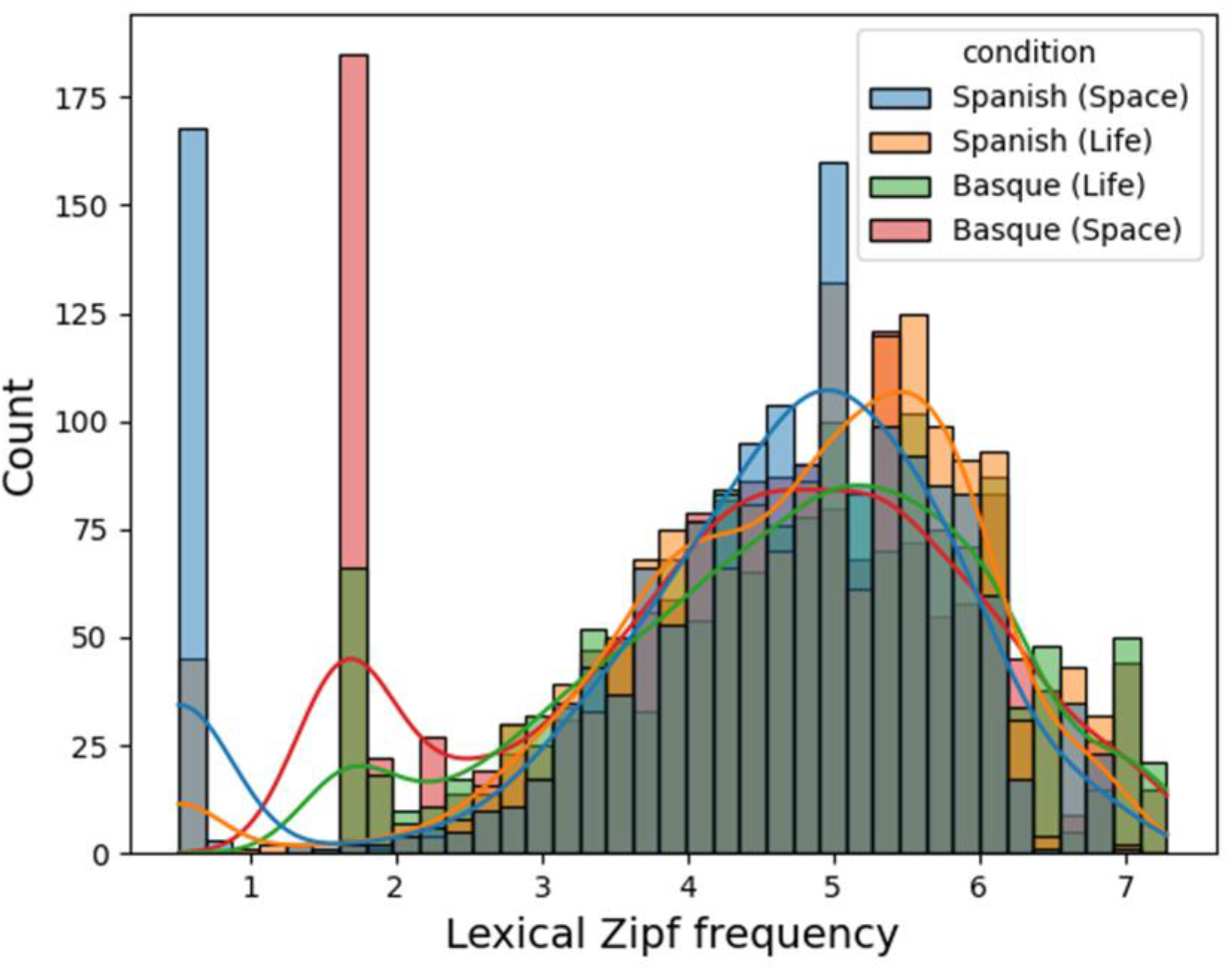
Zipf frequency distributions of content words in Spanish (1st, blue; 2nd, yellow), and Basque (1st, green; 2nd, red) stories. Bars represent content word count per Zipf frequency bin and smooth lines mark the density distribution.

**Supplemental Figure 2.**
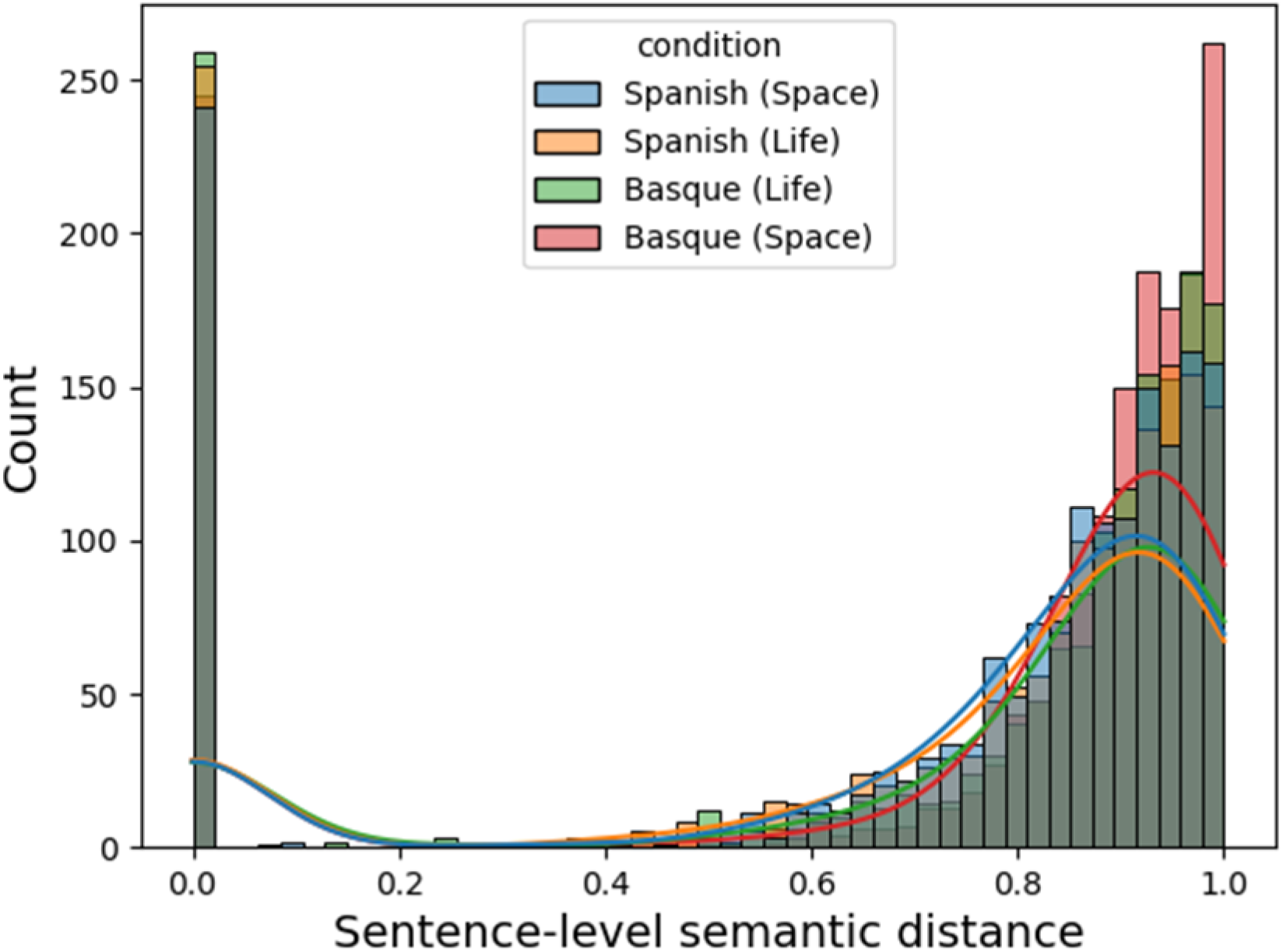
Sentence-level semantic distance distributions of content words in Spanish (1st, blue; 2nd, yellow), and Basque (1st, green; 2nd, red) stories. Bars represent content word count per bin of sentence-level semantic distance and smooth lines mark the density distribution.

**Supplemental Figure 3.**
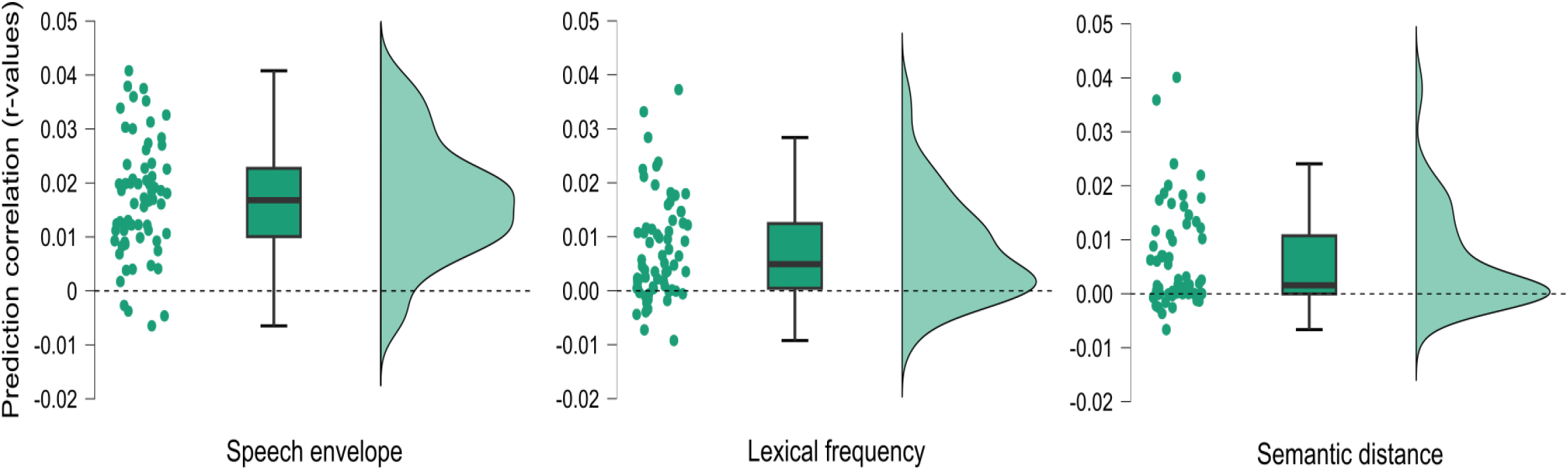
Distributions of average prediction correlations between each regressor and the mTRF model. Each participant is represented by two points (one per language). In the boxplots, horizontal lines within each box mark the median score. Upper and lower hinges mark the first and third quartile, and whiskers show 1.5 * interquartile range. Density plots mark the group-level distribution of prediction correlations for each regressor.

**Supplemental Formula 1.** Formula and rationale of the amount of exposure (AoE) to a language composite index (*AoE_L_*). The *CurrentHomeAoE_L_* and *CurrentSchoolAoE_L_* values are themselves composite indexes (*SF2* and *SF3* respectively):

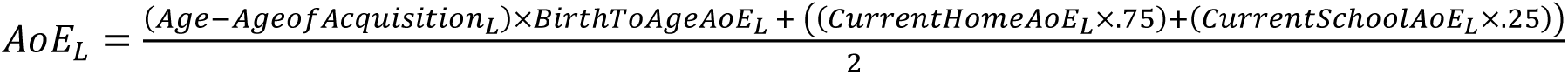

The rationale behind the multipliers of the different indexes that build the composite measure is weighting the contexts in which participants are exposed to a given language by the relative number of waking hours that they represent in an everyday life situation. We took into consideration the relative amount of waking hours that children spend at home (∼3147 h; 75% of the total of waking hours) and at school (∼1050 h; 25% of waking hours), as well as the relative amount of school hours that children spend in classes (∼875 h; 83%) and leisure time (∼175 h; 17%) according to their schooling system (Official Gazette of the Basque Country, 2008).

**Supplemental Formula 2.** Formula of the *CurrentHomeAoE_L_* composite index:

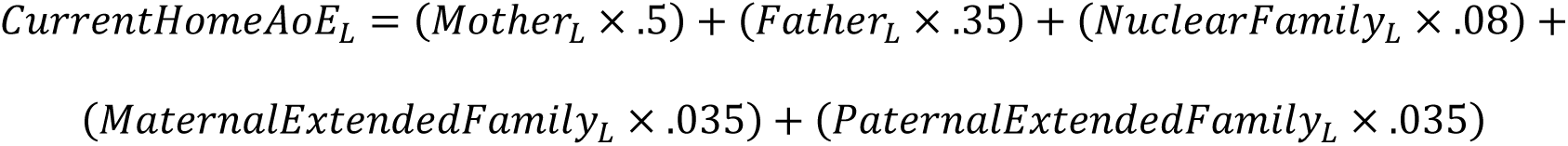

In Supplemental Formula 2, we took into consideration the proportion of waking hours that children spend in contact with the different members of their familial environment (Leaper et al., 1998; Pancsofar & Vernon-Feagans, 2006). In the case of one-parent families, we adapted the above formula to *CurrentHomeAoE_L_*\*, which aggregates the input from both parents into a single value (*Parent_L_*), as well as the ones from the maternal and paternal extended family (*Extended Family_L_*), as follows:
**Supplemental Formula 3.** Formula of the CurrentSchoolAoEL composite index:

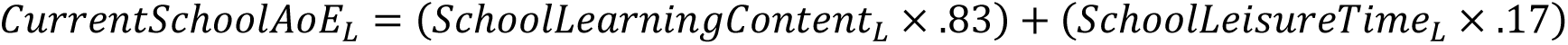

